# Ancestral Tributyltin Exposure Produces Male-Specific Brown Adipose Thermogenic Dysregulation

**DOI:** 10.1101/2025.11.04.686561

**Authors:** Yikai Huang, Richard C. Chang, Katelyn Uyen Nguyen, Kaitlin To, Myah Gibbs, Harunori Honjoh, Toshi Shioda, Bruce Blumberg

## Abstract

Obesity is a global health challenge increasingly linked to environmental obesogens. Tributyltin (TBT), a model obesogen that activates PPARγ and RXR, promotes transgenerational obesity in mice, but whether developmental and ancestral exposure impairs energy balance remains unclear. Brown adipose tissue (BAT) contributes to energy expenditure and thermoregulation through UCP1-dependent and -independent pathways. We tested whether TBT exposure disrupts BAT function in F1 and F3 descendants of exposed F0 dams.

We measured body composition before and after high-fat diet (HFD) challenge and assessed resting core body temperature, BAT gene expression, mitochondrial DNA (mtDNA) content, and mitochondrial electron transport chain (ETC) capacity. TBT-line males accumulated more fat after HFD without changes in lean mass despite similar mass-scaled food intake. Before HFD, core body temperature was lower in F1 and F3 TBT-line males, indicating impaired thermal homeostasis before increased adiposity. Candidate-gene qPCR did not show uniform suppression of thermogenic genes; instead, Ucp1 mRNA increased while Ucp2 mRNA decreased in F1 males, indicating dysregulation. Transcriptomal profiling revealed altered thermogenic signaling and consistent downregulation of mitochondrial electron transport chain genes across F1 and F3 males. Relative mtDNA content was unchanged, whereas complex-specific respirometry revealed reduced complex I- and II-associated respiratory capacity, providing evidence of impaired electron transport. No consistent effects occurred in females.

These findings identified an early, male-associated BAT phenotype characterized by thermogenic dysregulation, reduced mitochondrial ETC capacity, and impaired thermal homeostasis. They support a model in which impaired BAT mitochondrial function contributes to later susceptibility to diet-induced adiposity.

## Introduction

Obesity has increased worldwide over the past several decades and is now a major risk factor for diabetes, cardiovascular disease, and other chronic illnesses [1, 2]. While excessive caloric intake and reduced physical activity are important contributors, these lifestyle factors do not fully explain the rapid rise in obesity prevalence [3, 4]. Environmental chemicals with endocrine-disrupting activity, termed obesogens, have emerged as additional contributors [5, 6]. Obesogens can alter developmental programs, promote adipogenesis, and increase susceptibility to obesity later in life [7, 8].

Tributyltin (TBT) is a well-studied obesogen that activates the nuclear hormone receptors PPARγ and RXR to promote adipocyte differentiation [9, 10]. Previous studies showed that TBT exposure of pregnant F0 dams increased adiposity in directly exposed F1 and F2 animals and in their unexposed F3 and F4 descendants, indicating transgenerational effects [11–14]. These findings suggest that obesogen exposure can leave a heritable mark on metabolism. However, the mechanisms linking TBT exposure to long-term metabolic outcomes are not fully understood.

Brown adipose tissue (BAT) is a key regulator of energy balance because it oxidizes metabolic substrates to generate heat and contributes to whole-body energy expenditure and metabolic rate [15–17]. Classical non-shivering thermogenesis is mediated by uncoupling protein 1 (UCP1), an inner mitochondrial membrane protein that dissipates the mitochondrial proton-motive force as heat rather than coupling it to ATP synthesis in BAT [15, 18]. UCP1-dependent thermogenesis requires appropriate UCP1 expression and activation but also sufficient mitochondrial respiratory capacity to sustain substrate oxidation, electron transport, and proton-gradient formation. Consistent with this functional dependence, disruptions in respiratory-chain composition or mitochondrial DNA (mtDNA)-encoded electron transport chain (ETC) subunits impaired BAT mitochondrial respiration and thermogenic function [19]. Thus, defects in mitochondrial ETC capacity may compromise BAT energy expenditure even when changes in *Ucp1* expression alone do not fully explain the thermogenic phenotype.

BAT also acts as an endocrine organ by secreting batokines such as FGF21, IL-6, and 12,13-diHOME, which regulate glucose and lipid metabolism and contribute to systemic energy balance [20]. Impaired BAT function may affect whole-body energy balance by decreasing thermogenesis and altering systemic metabolic signaling. Human studies support the physiological relevance of BAT, but they also highlight important caveats. Most active BAT depots in adult humans resemble beige adipocytes rather than classical brown adipocytes [16, 21, 22], and adrenergic signaling differs between human and mice; β2-AR predominates in humans versus β3-AR in mice [21, 22]. Environmental context is also critical. Housing mice at sub-thermoneutral temperatures (∼22 °C) chronically activates BAT, whereas many modern humans spend substantial time in climate-controlled environments that reduce chronic cold exposure and BAT activation [16, 21]. Intriguingly, population studies suggest that average basal metabolic rate (BMR) has declined over the past ∼100 years, with warmer ambient environments and reduced BAT activation proposed as potential contributing factors [23, 24]. Together, these observations motivate assessing whether changes in BAT are accompanied by a measurable difference in thermal homeostasis in mice, alongside direct analysis of BAT function.

Here, we examined whether developmental and ancestral TBT exposure disrupted BAT function in F1 and F3 descendants before HFD challenge. We evaluated body composition, resting core body temperature, a predefined panel of BAT identity and thermogenic genes, broader BAT transcriptomic profiles, relative mtDNA content, and mitochondrial ETC function. Our results showed that TBT-line males accumulated more fat mass after HFD without changes in lean mass despite similar mass-scaled food intake. TBT line males also exhibited lower resting core body temperature, mixed changes in targeted thermogenic gene expression, consistent transcriptional suppression of mitochondrial ETC-related genes, and reduced complex-specific mitochondrial respiratory capacity without a detectable change in relative mtDNA content before obesity development. These findings support a model in which early BAT mitochondrial dysfunction contributes to impaired thermal homeostasis and increases the susceptibility of TBT-line males to diet-induced obesity.

## Material and Method

### Chemicals and Reagents

Tributyltin chloride (TBT) was obtained from Sigma-Aldrich (St. Louis, MO). Carboxymethyl cellulose (CMC), dimethyl sulfoxide (DMSO), and other standard reagents were purchased from Sigma-Aldrich unless otherwise specified. RNA isolation reagents (Trizol, Thermo Fisher Scientific), cDNA synthesis kits (High-Capacity cDNA Reverse Transcription Kit, Applied Biosystems), and qPCR reagents (SYBR Green PCR Master Mix, Thermo Fisher Scientific) were used according to manufacturer protocols.

### Animal Maintenance and Exposure

C57BL/6J mice were obtained from The Jackson Laboratory (Sacramento, CA) and maintained in microisolator cages at the University of California, Irvine animal facility under controlled environmental conditions (23–24 °C, 12 h light/dark cycle) with ad libitum access to food and water. All animal experiments were approved by the Institutional Animal Care and Use Committee of the University of California, Irvine and performed in a manner consistent with Federal guidelines.

For these transgenerational experiments designated transgenerational experiment 5 (T5) and transgenerational experiment 7 (T7), female mice were exposed to 50 nM TBT or 0.1% DMSO vehicle (both diluted in 0.5% CMC) via drinking water from preconception throughout pregnancy and lactation, as previously described[11, 12]. The selected concentration was based on prior studies and is five-fold lower than the established no observed adverse effect level (NOAEL) for rodents [25]. Only litters with 6-8 pups and at least two of each sex were selected to minimize litter size effects on metabolic endpoints [26] as in our previous studies. To avoid pseudoreplication, only one male and one female per litter were selected for endpoint analyses, while additional siblings were used for breeding to produce the next generation [26]. Randomized breeding was performed across litters, and siblings were never mated with each other.

### Diet Challenge and Body Composition Analysis

From weaning until 12 weeks of age, all offspring were maintained on a chow diet (PicoLab 5053, 24.5% kcal from protein, 13.1% kcal from fat, 62.3% kcal from carbohydrate). At 12 weeks, mice were switched to a higher fat diet (PicoLab 5058, 20.3% kcal from protein, 21.6% kcal from fat, 58.1% kcal from carbohydrate) for 6 weeks. Body fat and lean mass were measured weekly from 12 weeks (baseline) to 18 weeks (endpoint) using EchoMRI™ Whole Body Composition Analyzer, as previously described [11, 12, 27]. Food intake was monitored twice weekly. Kleiber’s law analysis was conducted by normalizing food intake to body mass^0.75 [28, 29].

### Core Body Temperature Measurement

Core body temperature was measured at 12 weeks of age, before HFD challenge and at 18 weeks of age, after HFD challenge in F1 F2 and F3 TBT-line and control mice from an independent cohort, designated as transgenerational experiment 7 (T7). Resting rectal temperature was measured at approximately 8:00 a.m. using a probe (RET-3 Rectal Probe for Mice, Physitemp, LLC) connected to a thermometer (Therma Waterproof thermometer, ThermoWorks^©^). The probe was inserted to a consistent depth of 1 cm, and the temperature was recorded after the reading stabilized. One measurement was collected from each mouse under the same environmental conditions. Data was analyzed using GraphPad Prism.

### Tissue Collection

At 12 weeks (pre-diet) and 18 weeks (post-diet), mice were euthanized by CO₂ asphyxiation followed by cervical dislocation. Interscapular BAT was dissected, weighed, and snap-frozen in liquid nitrogen for RNA/DNA extraction or functional assays.

### Quantitative Real-Time PCR (qPCR)

Total RNA was extracted from BAT using Trizol and reverse transcribed with the High-Capacity cDNA Reverse Transcription Kit (Applied Biosystems). qPCR was performed with SYBR Green PCR Master Mix on a Roche LightCycler 480 II system. Cycle threshold values were determined using the second derivative maximum method, and relative expression was calculated by the 2^−ΔΔCt^ method [15], adjusted for primer efficiency [30]. Standard propagation of error was used throughout each treatment group [31, 32]. Error bars represent the standard error of the mean (SEM) from 10 to 12 biological replicates, calculated using standard propagation of error. Expression was normalized to *Hprt*. Primer sequences and efficiencies are listed in Supplementary Table S1[33].

### RNA sequencing

Total RNA was extracted from BAT samples using the Direct-zol™ RNA MiniPrep Kit (Zymo Research) according to the manufacturer’s instructions. RNA-sequencing libraries were subsequently constructed from qualified RNA samples and sequenced on Illumina NextSeq2000 sequencer. Raw sequencing reads underwent quality-control assessment and were aligned to the mouse reference genome (GRCm38/mm10) using STAR [34]. Gene-level read counts were generated using featureCounts [35], and differential expression analysis was performed using DESeq2 [36, 37]. RNA sequencing datasets are accessible through Gene Expression Omnibus (GEO Series GSE345574).

### mtDNA Copy Number Assay

Genomic DNA was isolated from BAT using the Zymo D3025 kit. Relative mtDNA copy number was determined by qPCR using primers targeting mitochondrial (*Nd1* and *16SrRNA*) genes, as described previously in Quantitative Real-Time PCR (qPCR) of Method section [38]. Ratios of mitochondrial to nuclear amplicons were calculated to estimate relative mtDNA content.

### Seahorse Analysis of BAT Respiration

Frozen stored BAT tissue from 12-(pre-diet) and 18-week (post-diet)-old F1 and F3 mice were used to isolate mitochondria. Isolated mitochondrial extracts were quantified by Qubit™ Protein and Protein Broad Range (BR) Assay Kits. Oxygen consumption rate (OCR) was measured using a Seahorse XFe96 Analyzer (Agilent Technologies). Mitochondria extracts were restored for ETCP with mixture of Alamethicin, NADH, Cyt C with Mitochondrial Respirate Buffer with BSA. Complex 1, 2 and 4 respiration capacity were tested following sequential injections of rotenone, succinate; antimycin A; ascorbate, TMPD and sodium azide. OCR values were normalized to mitochondria mass [39, 40]. Complex I–associated OCR was calculated as the NADH-supported OCR measured before rotenone/succinate injection minus the antimycin A–insensitive OCR; complex II–associated OCR was calculated as the succinate-supported OCR measured after rotenone/succinate injection minus the antimycin A–insensitive OCR; and complex IV–associated OCR was calculated as the ascorbate/TMPD-supported OCR minus the sodium azide–insensitive OCR [41].

### Statistical Analysis

All data were analyzed using GraphPad Prism 10. Statistical comparisons were performed using two-way ANOVA (factors: treatment and diet) with Tukey’s post-hoc test. Data are presented as mean ± SEM. Statistical significance was defined as p < 0.05.

## Results

### TBT-line males accumulated more fat after dietary challenge despite unchanged lean mass and similar mass-scaled food intake

To determine whether ancestral TBT exposure altered whole-body composition, we measured fat (Figure 1A) and lean mass (Figure 1B) in F1 and F3 males at 12 weeks of age (pre-diet) and after a 6-week HFD challenge (18 weeks). We found no difference in fat or lean mass between groups at baseline. However, after HFD challenge, TBT-line males accumulated substantially more fat mass than controls whereas lean mass remained unchanged. To assess whether these changes reflect altered intake or efficiency, we analyzed food intake using Kleiber’s law scaling (Figure 1C). Normalized food intake was similar between groups despite the greater fat accumulation in TBT-line males. Thus, the increased adiposity was not accompanied by a detectable increase in food intake and may instead involve altered energy expenditure, metabolic efficiency, or nutrient partitioning [29, 39]. No differences were observed in females (Figure 1D-F). We also weighed BAT at tissue collection and expressed its weight relative to body weight (Supplementary Figure S1A–H[33]). Relative BAT weight was lower in pre-diet F1 TBT-line males than in controls, while no significant lineage differences were detected in the other groups.

**Figure 1.**
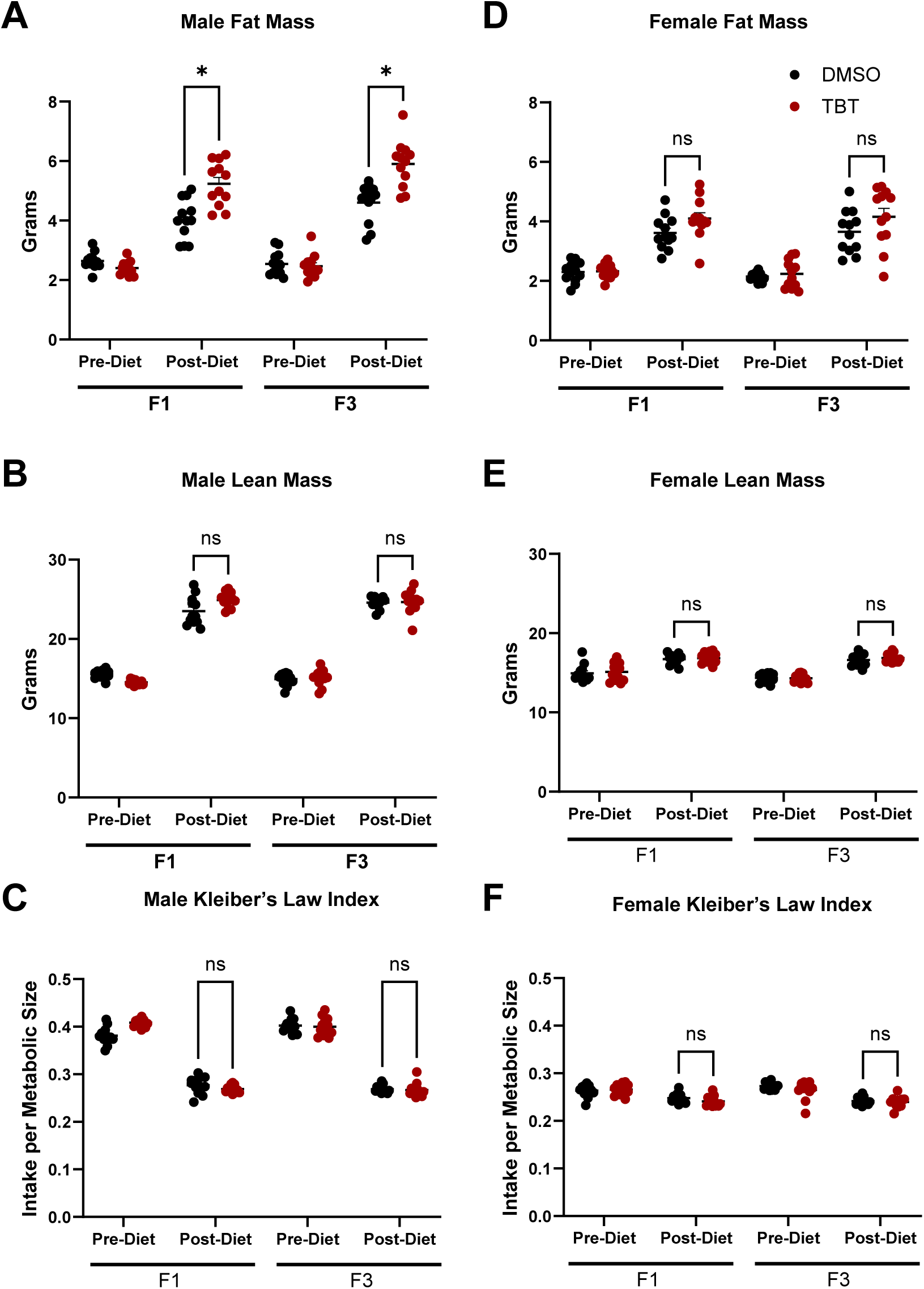
Ancestral TBT exposure increases fat mass in male offspring without altering lean mass or food intake. (A–C) Body composition and food intake of F1 and F3 male offspring. Fat mass (A) and lean mass (B) were measured by EchoMRI at 12 weeks (Pre-Diet) and after a 6-week high-fat diet (Post-Diet). (C) Food intake normalized to body mass^0.75 (Kleiber’s law index) was calculated to assess energy intake relative to metabolic size.(D–F) Corresponding data for female offspring. TBT-line males exhibited significantly higher fat mass after high-fat diet challenge in both F1 and F3 generations, with no difference in lean mass or normalized food intake. Female offspring showed no differences across groups.Data are presented as mean ± SEM; each point represents one animal (one male or female per litter; n = 12). Two-way ANOVA (factors: treatment, diet) with Tukey’s post-hoc test; *p < 0.05, ns = not significant.

### TBT-line males exhibited lower resting core body temperature before dietary challenge

To determine whether TBT-line males exhibited an early physiological phenotype before the development of HFD-induced obesity, we measured resting core body temperature using a rectal temperature probe in an independent T7 cohort generated under the same exposure, breeding, dietary, and age conditions as the cohort used for the other analyses (Figure 2). Under control-diet (CD) conditions at 12 weeks of age, TBT-line males exhibited significantly lower rectal temperature than DMSO-control males in the F1, F2, and F3 generations (Figure 2A–C). After 6 weeks of HFD feeding, both DMSO-control and TBT-line males exhibited lower mean body temperatures than under control-diet (CD) conditions, and the difference between lineages was no longer significant in any generation. No significant differences were detected between TBT-line and DMSO-control females under either dietary condition in any generation (Figure 2D–F). Together, these findings demonstrate a male-specific reduction in resting core body temperature before HFD challenge and a distinct post-HFD pattern in which both lineages exhibited lower mean temperatures without a detectable between-lineage difference, consistent with poorer maintenance of core body temperature following HFD exposure.

**Figure 2.**
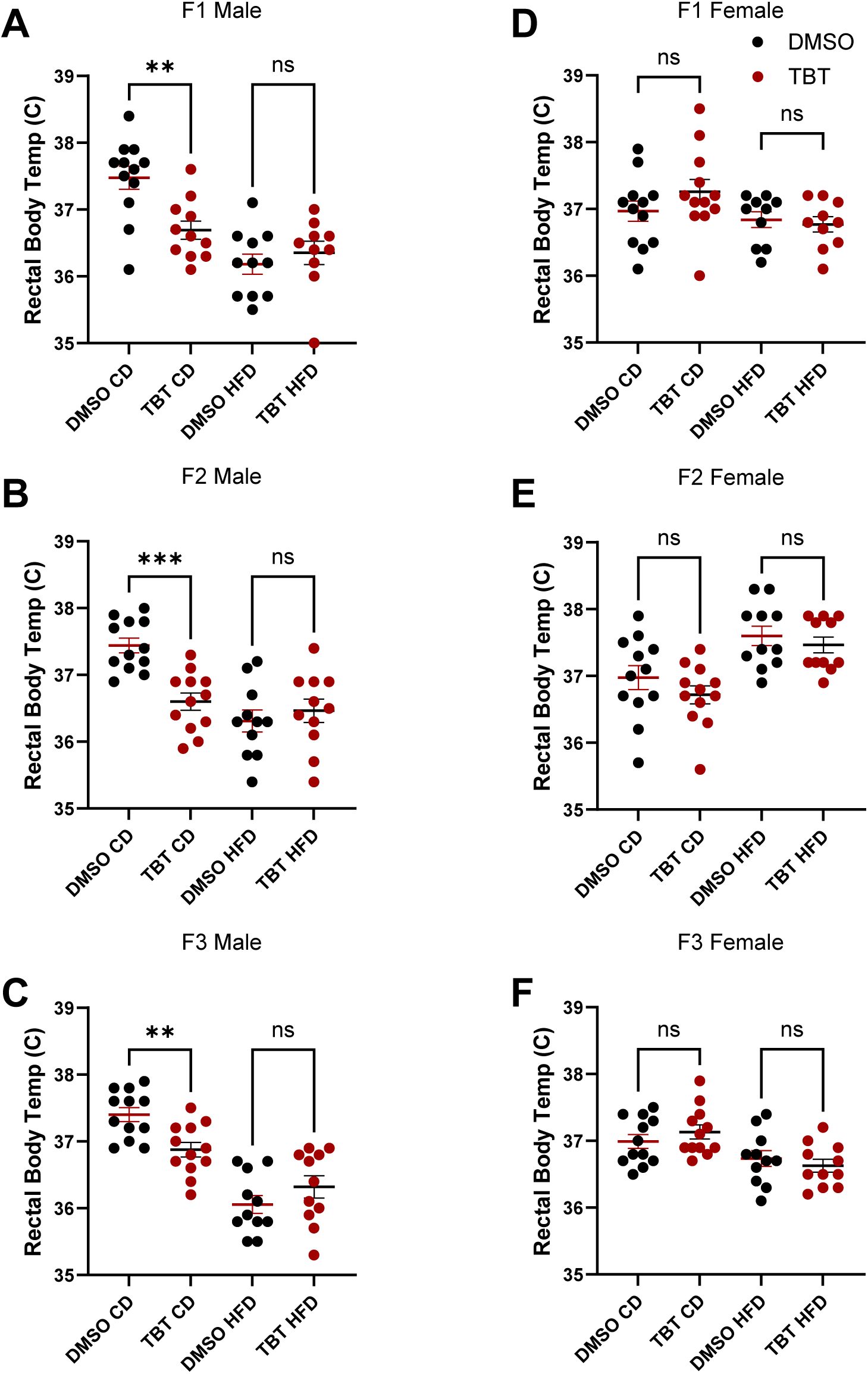
Rectal core body temperature in male and female offspring. Rectal body temperature was measured in F1, F2, and F3 male (A–C) and female (D–F) offspring from DMSO- or TBT-exposed lineages at 12 weeks of age while maintained on control diet (CD; pre-diet) and at 18 weeks of age after 6 weeks of high-fat-diet (HFD) feeding. Each point represents an individual mouse (n = 12 per group); horizontal red bars indicate mean ± SEM. Statistical significance was determined using two-tailed Student’s t-tests comparing DMSO and TBT groups within each generation and dietary condition. **P < 0.01; ***P < 0.001; ns, not significant. Under CD conditions at week 12, TBT-line males exhibited significantly lower core body temperature in all three generations. This difference was no longer detected after HFD feeding at week 18. No significant differences were detected in females under either dietary condition. These findings identify a male-specific reduction in basal thermal homeostasis before HFD challenge that was not maintained after HFD feeding.

### TBT-line males BAT showed potential dysregulated thermogenesis genes expression

To determine whether the BAT gene-expression pattern we previously identified in obese TBT-line males [42] was already present before HFD challenge, we measured BAT identity-associated marker genes (*Zic1, Ebf2, Pdk4*) (Figure 3 A-C) and some key thermogenic marker gene (*Pgc1α, Ucp1, Ucp2*) expression in F1 and F3 pre-diet males (Figure 3D-F). The pre-diet expression pattern did not show uniform suppression of the tested genes. *Zic1* and *Ebf2* were not significantly altered in either generation, although *Ebf2* mRNA levels were lower in both generations and *Zic1* mRNA levels were lower in F3 males. *Pdk4* was significantly reduced in F3 males but was unchanged in F1 males. Similarly, *Pgc1a* mRNA levels were not significantly altered in either generation. However, in F1 males *Ucp1* mRNA levels were significantly increased, whereas *Ucp2* mRNA levels were significantly decreased. mRNA levels were not significantly altered in F3 males. These opposing changes in *Ucp1* and *Ucp2* mRNA levels, together with the generation-specific reduction in *Pdk4* mRNA levels, is consistent with dysregulation rather than uniform suppression of the thermogenic gene-expression program before HFD challenge.

**Figure 3.**
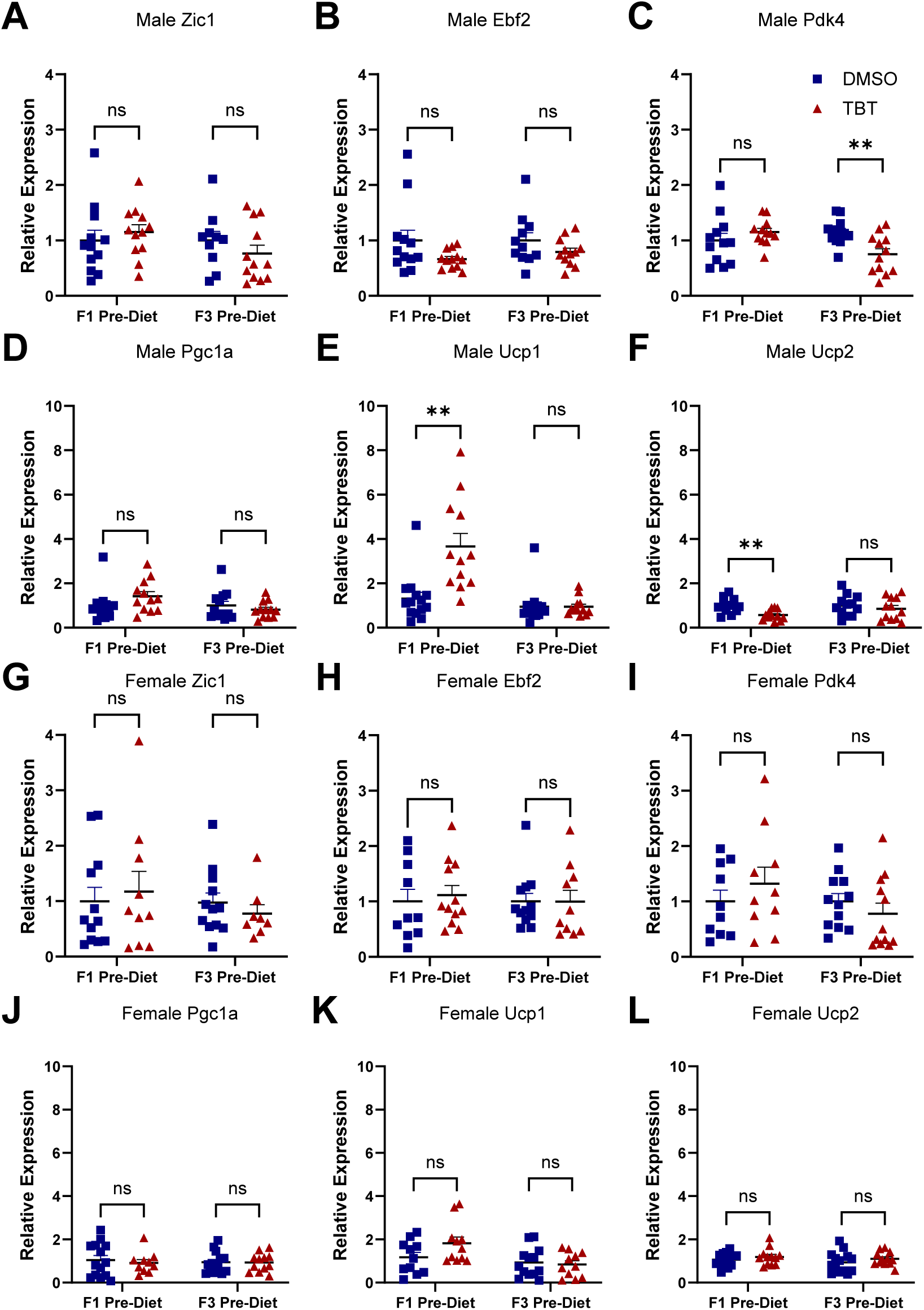
Expression of BAT identity-associated and thermogenic genes in pre-diet offspring. (A–C) Expression of brown adipose identity marker genes Zic1, Ebf2, and Pdk4 in BAT from F1 and F3 male offspring measured by qPCR at 12 weeks (Pre-Diet). (D–F) Expression of thermogenesis marker genes Pgc1a, Ucp1, and Ucp2 in the corresponding male offspring. (G–I) Expression of Zic1, Ebf2, and Pdk4 in F1 and F3 female offspring measured by qPCR at 12 weeks (Pre-Diet). (J–L) Expression of Pgc1a, Ucp1, and Ucp2 in the corresponding female offspring. Data are mean ± SEM; each point represents one animal (one male or female per litter; n = 12). Two-way ANOVA (factors: treatment and diet) with Tukey’s post-hoc test; *p < 0.05, **p < 0.01, ***p < 0.001; ns, not significant.

For comparison, after 6 weeks of HFD feeding, *Zic1* and *Ebf2* mRNA levels were significantly reduced in both F1 and F3 TBT-line males (Supplementary Figure S2A–B[33]). *Pdk4* showed generation-dependent changes, with increased mRNA levels in F1 males and decreased mRNA levels in F3 males (Supplementary Figure S2C[33]). *Pgc1a* mRNA levels were increased in F1 males but not significantly altered in F3 males, whereas *Ucp2* mRNA levels were decreased only in F1 males and *Ucp1* mRNA levels were not significantly altered in either generation (Supplementary Figure S2D–F[33]). Thus, the post-diet pattern showed reduction in mRNA levels for *Zic1* and *Ebf2* but remained gene- and generation-dependent.

No significant differences in mRNA levels for the tested genes were detected in pre-diet females (Figure 3G–L). After HFD challenge, *Ucp1* mRNA levels were increased in F1 TBT-line females, whereas the other measured genes were not significantly altered in either generation (Supplementary Figure S2G–L[33]). Together, these results indicate a potential thermogenic dysregulation may exist in BAT in TBT-line males even before the development of obesity was triggered [39, 40, 43]. However, the mixed and generation-dependent changes detected by the targeted qPCR panel did not reveal a sufficiently consistent molecular pattern to explain the early physiological phenotype.

### Transcriptomal profiles of TBT-line males BAT demonstrated dysregulated thermogenic genes

To address potential changes in gene expression patterns related to thermogenic activities in non-obese pre-diet mice, we performed bulk RNA sequencing. Differential expression analysis identified significant changes in both thermogenesis-related and electron transport chain/oxidative phosphorylation (ETC/OXPHOS)-related genes in pre-diet TBT-line males compared with DMSO controls (Figure 4A–D). The F1 male comparisons showed extensive bidirectional changes in both gene sets, whereas fewer genes were significantly altered in F3 pre-diet males. Analysis of thermogenesis-related transcriptional programs showed that mRNAs encoding several upstream signaling and regulatory genes were lower in TBT-line pre-diet males (Figure 4E). These changes included genes associated with natriuretic peptide–cGMP–PKG signaling (*Npr1* and *Prkg1*), the PKA–RSK–CREB signaling axis (*Creb1* and *Rps6ka2*), thermogenic transcription and BAT identity (*Foxc2, Ppargc1b, Kdm6b, Zfp516,* and *Smarca2*), MAPK–p38 signaling (*Map2k6*), and β-adrenergic receptor/cAMP generation (*Adcy7* and *Adcy1*). Downstream thermogenic regulators and effectors, however, were also not uniformly downregulated. For example, *Ppargc1a* and *Ucp1* mRNA levels were elevated in F1 TBT-line pre-diet males (Figure 4E). Thus, the lower mRNA levels for several upstream regulatory genes, together with inconsistent changes in mRNAs for downstream genes, indicated dysregulation of the thermogenic transcriptional program.

**Figure 4.**
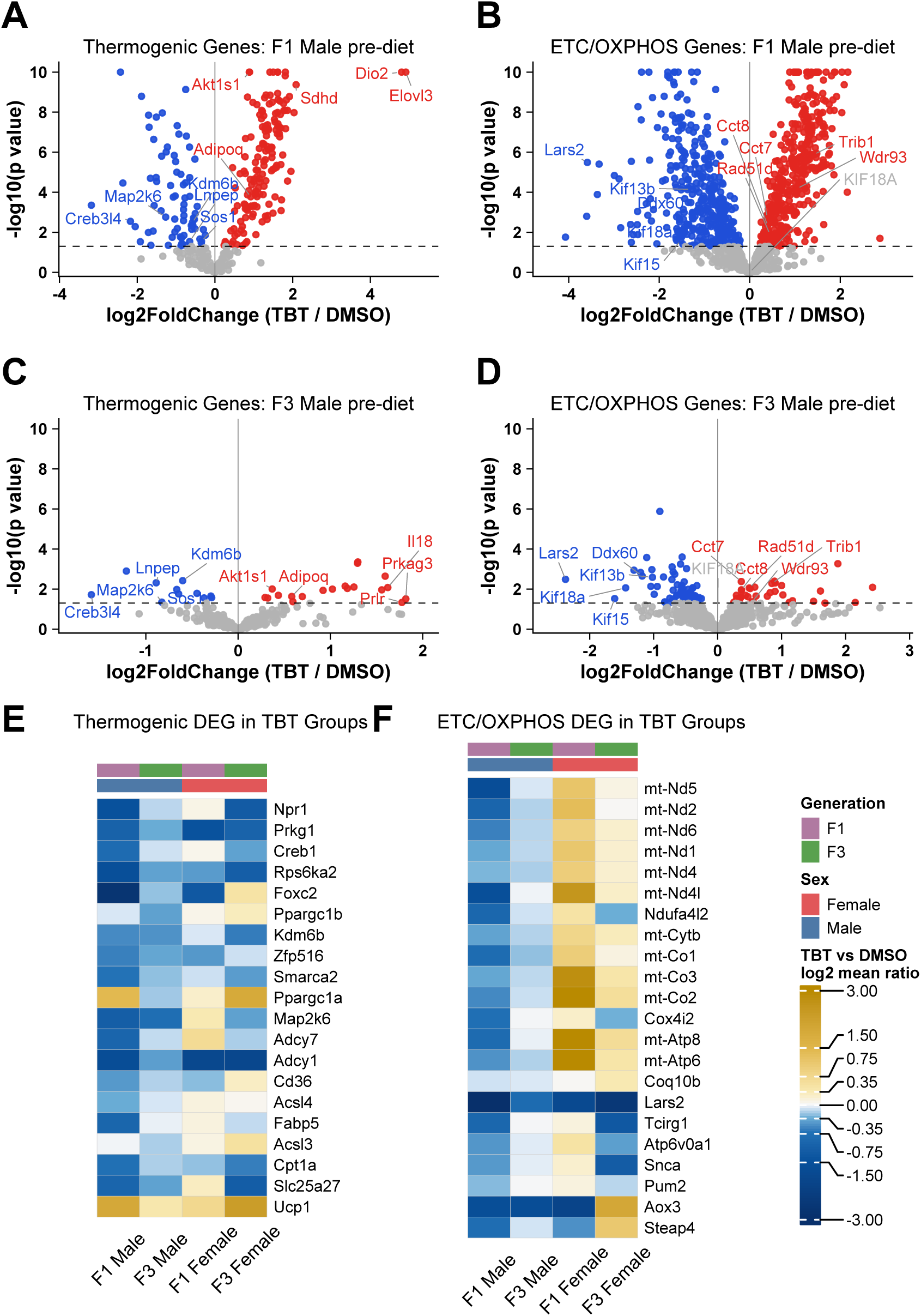
Transcriptomic dysregulation of thermogenic and mitochondrial respiratory-chain programs in pre-diet BAT. Bulk RNA sequencing was performed on brown adipose tissue (BAT) collected from F1 and F3 offspring at 12 weeks of age before high-fat diet challenge. (A, C) Volcano plots of genes associated with the thermogenesis pathway in F1 (A) and F3 (C) TBT-line males compared with their corresponding DMSO controls. (B, D) Corresponding volcano plots of electron transport chain and oxidative phosphorylation (ETC/OXPHOS)-related genes in F1 (B) and F3 (D) males. Each point represents one gene. The x-axis shows the log2 fold change in TBT-line relative to DMSO control BAT, and the y-axis shows −log10(p value). Significantly increased and decreased genes in TBT-line BAT are shown in red and blue, respectively, while nonsignificant genes are shown in gray. Top 10 changed genes are labeled, and the horizontal dashed line indicates p = 0.05. (E) Heatmap showing the TBT-to-DMSO log2 mean expression ratio of selected thermogenesis-related genes across F1 male, F3 male, F1 female, and F3 female BAT. Genes are organized by thermogenesis-related signaling and functional programs. (F) Heatmap showing the corresponding log2 mean expression ratios of selected ETC/OXPHOS-related genes organized by respiratory-chain complex and functional category. Blue indicates lower expression in TBT-line BAT, white indicates little or no change, and gold indicates higher expression relative to DMSO controls. Thermogenesis-related genes showed mixed and sex- and generation-dependent changes, consistent with dysregulation rather than uniform suppression of the thermogenic program. Selected ETC/OXPHOS-related genes showed recurrent decreases in both F1 and F3 males, whereas changes in females were more variable and gene-specific.

The heatmap summarized ETC/OXPHOS-related genes that showed lower mRNA levels in both F1 and F3 TBT-line pre-diet males (Figure 4F). These genes included mtDNA-encoded subunits of respiratory complexes I (*mt-Nd5, mt-Nd2, mt-Nd6, mt-Nd1, mt-Nd4,* and *mt-Nd4l*), III (*mt-Cytb*), IV (*mt-Co1, mt-Co2,* and *mt-Co3*), and V (*mt-Atp6* and *mt-Atp8*), as well as nuclear-encoded subunits (*Ndufa4l2* and *Cox4i2*) and genes involved in the CoQ/ubiquinone pool, mitochondrial translation, and respiratory-chain support (*Coq10b, Lars2, Tcirg1, Atp6v0a1, Snca, Pum2, Aox3,* and *Steap4*). After HFD challenge, thermogenesis- and ETC/OXPHOS-related genes remained differentially expressed in male BAT, but changes across the same functional panels were mixed and generation-dependent, and the recurrent pre-diet decreases were not uniformly maintained (Supplementary Figure S3A–F[33]). Female BAT also showed significant transcriptional changes in these pathways, but the direction of change varied by generation and dietary condition; notably, mRNA levels for several mtDNA-encoded respiratory-complex genes were increased in F1 pre-diet females, contrasting with the recurrent decreases observed in pre-diet males (Figure 4E–F; Supplementary Figure S4A–H[33]). Together, these results revealed broader transcriptional dysregulation of thermogenic and mitochondrial respiratory-chain programs than was detected by the targeted qPCR panel and prompted us to assess mitochondrial abundance and respiratory-chain capacity directly.

### Reduced complex-specific respiratory activity occurred without a detectable change in relative BAT mtDNA content

To determine whether altered gene expression reflected reduced mitochondrial abundance, we measured mtDNA copy number in F1 and F3 males at 12 weeks of age before HFD challenge (Figure 5A-B). We found that mtDNA copy number was not changed at baseline in TBT-line males, which indicates the transcriptomal changes in TBT-line male or potential organ dysfunction were not the result of altered mtDNA abundance. In females, ND1 copy number was unchanged in both generations, although 16S copy number was increased in F1 TBT-line females but unchanged in F3 females (Supplementary Figure S5A–B[33]), indicating no consistent reduction in relative BAT mtDNA content. To directly assess function, we performed Seahorse assays on frozen BAT mitochondrial extracts. Seahorse-based assessments are widely used to evaluate BAT oxidative capacity and provide functional readouts of mitochondrial energetics [39, 40]. Modified ETCP Seahorse assessments made analyzing frozen tissue mitochondria complex capacity possible [41, 44]. OCR profiles were obtained following sequential addition of rotenone/succinate, antimycin A, ascorbate/TMPD, and sodium azide (Figure 5C–D). Complex I- and complex II-associated respiratory capacities were significantly reduced in both F1 and F3 TBT-line pre-diet males, whereas complex IV-associated respiratory capacity was unchanged (Figure 5E–J). Same ETCP Seahorse assay was performed on corelating female cohort samples as well (Supplementary Figure S5C-D[33]). No significant differences in complex I-, II-, or IV-associated respiratory capacity were detected in F1 or F3 females (Supplementary Figure S5E–J[33]). Together, these results showed that TBT-line males exhibited reduced complex I- and II-associated respiratory capacity before HFD challenge without a corresponding reduction in relative mtDNA content, indicating that the respiratory impairment reflected mitochondrial functional defects rather than reduced mitochondrial abundance.

**Figure 5.**
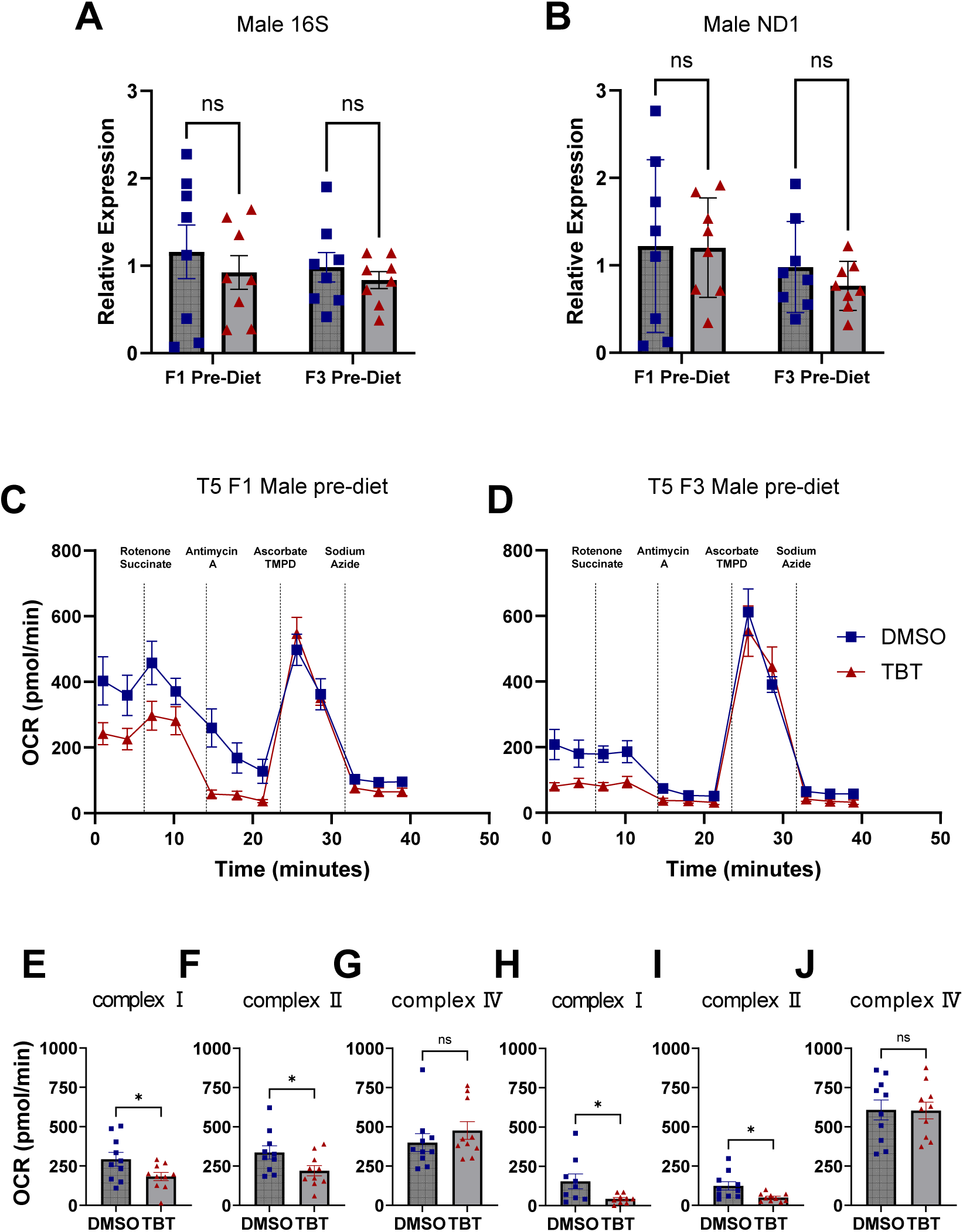
Relative mitochondrial DNA content and electron transport chain respiratory capacity in pre-diet BAT. (A–B) Relative mitochondrial 16S rRNA and ND1 copy numbers in brown adipose tissue (BAT) from F1 and F3 male offspring at 12 weeks (Pre-Diet). No consistent differences in relative mtDNA content were observed between DMSO- and TBT-line offspring. No significant differences were detected in males ND1 and 16S copy number. (C–D) Oxygen consumption rate (OCR) profiles obtained using the electron transport chain protocol (ETCP) in F1 and F3 male pre-diet BAT, respectively, following sequential addition of rotenone/succinate, antimycin A, ascorbate/TMPD, and sodium azide. (E–G) Quantification of complex I-, complex II-, and complex IV-associated respiratory capacity in F1 male BAT. (H–J) Corresponding quantification in F3 male BAT. TBT-line males exhibited reduced complex I and complex II respiratory capacity in both generations, whereas complex IV respiratory capacity was unchanged. All values are shown as mean ± SEM. Data were analyzed by unpaired t-test. *p < 0.05; ns, not significant.

## Discussion

We analyzed body composition, gene expression, mitochondrial respiratory function, and core body temperature in F1 and F3 males to assess whether developmental and ancestral TBT exposure impaired BAT function. We found that TBT-line males accumulated more fat mass after HFD challenge without changes in lean mass. Mass-scaled food intake was similar between groups, despite greater fat accumulation in TBT-line males. Relative BAT weight was significantly lower in pre-diet F1 TBT-line males than in DMSO controls, with no significant between-lineage differences in the other groups[33]. These data indicated that the increased adiposity was not accompanied by greater measured food intake and may instead involve altered energy expenditure, metabolic efficiency, or nutrient partitioning [15, 45].

To determine whether an early physiological phenotype was present before the development of HFD-induced obesity, we measured resting core body temperature in an independent cohort of F1, F2 and F3 male mice and female mice before and after HFD challenge. Before HFD challenge, resting core body temperature was lower in TBT-line males in all three generations, whereas no differences were detected in females. This pre-diet, male-specific reduction indicated impaired thermal homeostasis before obesity onset and raised the possibility that reduced thermogenic capacity contributed to the later susceptibility of TBT-line males to diet-induced obesity. After HFD feeding, both DMSO-control and TBT-line males showed lower mean body temperatures, and the difference between lineages was no longer significant, indicating that the post-diet response represented a distinct phenotype from the pre-diet TBT-associated deficit. This post-diet phenotype and its potential mechanisms were discussed in our previous work [42], and therefore are not discussed further here.

We next asked whether the disturbed pattern of gene expression levels we previously detected in BAT from obese TBT-line male mice [42] preceded obesity onset or was a consequence of it. We examined the same panel of marker genes in BAT from F1 and F3 pre-diet males. Notably, none of the tested BAT identity-related genes (*Zic1*, *Ebf2*, and *Pdk4*), which are associated with brown/beige adipocyte identity and metabolic regulation [46–49], was consistently altered in TBT-line males before HFD challenge. *Pgc1a*, another key regulator of mitochondrial biogenesis and thermogenic capacity, was likewise not significantly changed.

Although expression of several of these genes showed downward trends in at least one generation, the absence of significant and coordinated changes indicated that the expression pattern previously observed in obese TBT-line males was not fully present in the lean, pre-diet state. Interestingly, *Ucp1* and *Ucp2* exhibited opposing expression changes in F1 males, with *Ucp1* mRNA levels increased and *Ucp2* mRNA levels decreased. Neither gene showed significant changes in F3 males. This is consistent with dysregulation rather than uniform suppression of the tested thermogenic markers. Together, the discordant *Ucp1* and *Ucp2* expression changes raised the possibility that thermogenic dysregulation was already present in lean, pre-diet TBT-line males but involved genes or pathways not captured by the targeted panel tested. No significant changes in the tested genes were detected in pre-diet females. After HFD challenge, *Zic1* and *Ebf2* were significantly reduced in both generations of TBT-line males, while *Pdk4* was increased in F1 males but decreased in F3 males, *Pgc1a* was increased only in F1 males, *Ucp2* was decreased only in F1 males, and *Ucp1* remained unchanged in both generations. The female post-diet profile remained largely unchanged, except for increased *Ucp1* expression in F1 females. Thus, HFD challenge produced clearer suppression of specific BAT identity-related genes in males, although the broader targeted expression pattern remained gene- and generation-dependent.

Because the targeted panel did not reveal gene expression changes sufficient to explain the physiological phenotype, we used bulk RNA sequencing to establish a more comprehensive transcriptomal profile of pre-diet BAT from F1 and F3 TBT-line males. This analysis revealed a complex pattern of differential expression across thermogenesis-related pathways in both generations. Differentially expressed gene (DEG) analysis identified significant downregulation of genes involved in several thermogenesis-related pathways and functional programs, including the PKA–RSK–CREB signaling axis (*Creb1, Rps6ka1,* and *Rps6ka2*), brown-adipocyte and thermogenic regulation (*Foxc2, Kdm6b, Ppargc1b,* and *Zfp516*), and MAPK–p38 signaling (*Map2k6*). The discordance between the downregulation of several thermogenic regulatory genes and the increased expression of downstream thermogenic effector genes, including *Ucp1*, further indicated a loss of coordinated thermogenic gene regulation in TBT-line males before the development of HFD-induced obesity. DEG analysis also revealed a pattern of downregulation among genes involved in mitochondrial respiratory complex assembly and function in TBT-line males. After HFD challenge, thermogenesis- and ETC/OXPHOS-related genes remained differentially expressed in male BAT, but the direction and magnitude of these changes were mixed and generation-dependent, and the recurrent pre-diet decreases were not uniformly maintained. Female BAT also exhibited differential expression before and after HFD challenge, but the affected genes and directions of change varied by generation and dietary condition. Several mtDNA-encoded respiratory-chain genes showed higher mean expression in F1 pre-diet females, contrasting with the recurrent decreases observed in males. These results revealed broad transcriptional dysregulation across thermogenic pathways and directed our subsequent functional analysis toward mitochondrial function, particularly electron transport chain (ETC) respiratory capacity, in BAT from TBT-line males.

We measured relative mtDNA content and assessed complex-specific mitochondrial respiration to determine whether impaired mitochondrial function was accompanied by altered relative mtDNA content. Relative mtDNA content did not differ between TBT-line and control males in either the F1 or F3 generation, indicating that the respiratory defect was not accompanied by a detectable change in this indirect measure of mitochondrial content. In females, ND1 copy number was unchanged in both generations, although 16S rRNA copy number was increased in F1 TBT-line females but not in F3 females, indicating no consistent reduction in relative mtDNA content. In contrast, complex-specific respirometry revealed reduced ETC respiratory capacity in both generations, providing functional evidence of impaired mitochondrial function. Specifically, complex I- and II-associated respiratory capacities were reduced in F1 and F3 TBT-line males, whereas complex IV-associated respiratory capacity was unchanged. No significant differences in complex I-, II-, or IV-associated respiratory capacity were detected in females. Together, these findings indicate that reduced complex I and II respiratory capacity in TBT-line males occurred without a detectable change in relative mtDNA content, whereas corresponding respiratory defects were not detected in females.

Integrating the physiological, transcriptional, and mitochondrial findings, we propose a working model in which ancestral TBT exposure produces an early defect in BAT mitochondrial respiration. Reduced complex I and II respiratory capacities would limit electron entry into the electron transport chain. Because complex I contributes directly to proton pumping, whereas complex II supplies electrons to ubiquinone without pumping protons, impairment of both pathways could reduce downstream electron flow through complexes III and IV and weaken the electrochemical proton gradient, despite preserving intrinsic complex IV capacity. The resulting reduction in proton-motive force could constrain UCP1-mediated proton leak and heat production, thereby reducing BAT thermogenic capacity. Increased *Ucp1* expression may represent a compensatory response, although UCP1 protein abundance and activity were not measured. This impairment could contribute to reduced thermal homeostasis before dietary challenge and increase susceptibility to fat accumulation following HFD exposure.

Our findings add to a growing literature demonstrating that BAT is a systemic regulator of metabolic health. BAT secretes endocrine factors such as FGF21, IL-6, and 12,13-diHOME, which influence glucose and lipid homeostasis [20]. Therefore, impaired BAT function may affect whole-body energy balance by diminishing thermogenesis and altering systemic metabolic signaling. Recent human studies underscore the translational relevance of these mechanisms. Most active BAT depots in adults resemble beige rather than classical brown adipocytes [16, 21], and adrenergic signaling differs between species, with β2-adrenergic pathways predominating in humans compared to β3 in rodents [22]. Environmental context also matters. Mice are typically housed below thermoneutrality, which chronically activates BAT, whereas many modern humans spend substantial time in climate-controlled environments that reduce chronic cold exposure and BAT activation [21]. At the population level, average BMR has declined in recent decades, with warmer ambient environments and reduced BAT activation proposed as potential contributing factors [23, 24, 50, 51]. Together, these observations emphasize the importance of BAT in energy balance and the potential translational relevance of our findings.

Finally, our results should be viewed within the broader context of environmental obesogen research. Developmental exposure to DDT and DDE reduced sympathetic innervation and thermogenic gene expression in BAT [52], and other endocrine-disrupting chemicals such as BPA, phthalates, PFAS, and dioxins have been implicated in disrupting BAT or beige fat activity [53]. These observations suggest that BAT dysfunction may be a common target of obesogen action. To our knowledge, this study is the first to provide evidence that developmental and ancestral TBT exposure dysregulates BAT thermogenic programs and reduces mitochondrial respiratory capacity across generations.

Taken together, our findings show that TBT exposure produces transcriptional dysregulation of BAT thermogenic regulatory programs and reduced mitochondrial respiratory capacity in exposed F1 and unexposed F3 males. Whereas the post-HFD phenotype was characterized by altered broader thermogenic dysregulation and mitochondrial impairment were already present in lean, pre-diet TBT-line males. These transcriptomal and respiratory abnormalities occurred concurrently with lower resting core body temperature. This supports a model in which impaired mitochondrial respiration may constrain UCP1-dependent non-shivering thermogenesis and compromise BAT energy metabolism before obesity onset. The presence of related defects in F1 and F3 males supports both developmental and transgenerational consequences of TBT exposure but does not identify the underlying inheritance mechanism [11, 12, 42]. By integrating transcriptomal, mitochondrial, and physiological outcomes, our study supports BAT dysfunction as a candidate pathway linking developmental and ancestral obesogen exposure to later susceptibility to diet-induced obesity.

## Acknowledgements

This work was supported by grants from the National Institutes of Health, USA (R01ES023316 and R01ES031139) to B.B. We thank Dr. Dequina Nicholas (University of California) for providing access to the Seahorse instrument and for her advice on the design of the Seahorse experiments. We thank Dr. Munjal Acharya (University of California) and Dr. Hamendra Vekaria (Medical University of South Carolina) for their technical advice and helpful suggestions during the development of the Seahorse assay.

## Data Availability

The RNA-sequencing datasets generated and analyzed in this study are publicly available in the NCBI Gene Expression Omnibus (GEO) under accession number GSE345574. All other data generated or analyzed during this study are included in this article and its supplementary materials or are available from the corresponding author upon reasonable request.

---

**Supplementary Table S1. Primer information.** Primer sequences used for quantitative real-time PCR (qPCR) analyses. Forward and reverse primer sequences are listed in the 5′–3′ direction.

**Supplementary Figure S1. Relative BAT weight in male and female offspring before and after HFD challenge.** Brown adipose tissue (BAT) was weighed at tissue collection, and its weight was expressed as a percentage of body weight. Measurements were obtained from F1 males before HFD challenge (A) and after 6 weeks of HFD feeding (B), F3 males before (C) and after HFD (D), F1 females before (E) and after HFD (F), and F3 females before (G) and after HFD (H). Each point represents one mouse (n = 8 per group); horizontal red bars indicate mean ± SEM. DMSO and TBT groups were compared within each panel using an unpaired Welch’s *t* test. Relative BAT weight was significantly lower in pre-diet F1 TBT-line males than in DMSO controls; no significant differences were detected in the other groups. *P* < 0.05; ns, not significant.

**Supplementary Figure S2. Expression of BAT identity-associated and thermogenic genes in post-diet offspring.** (A–C) Expression of brown adipose identity marker genes *Zic1, Ebf2,* and *Pdk4* in brown adipose tissue (BAT) from F1 and F3 male offspring measured by qPCR at 18 weeks, after 6 weeks of high-fat diet feeding (Post-Diet). (D–F) Expression of thermogenesis marker genes *Pgc1a, Ucp1,* and *Ucp2* in the corresponding male offspring. (G–I) Expression of *Zic1, Ebf2,* and *Pdk4* in F1 and F3 female offspring. (J–L) Expression of *Pgc1a, Ucp1,* and *Ucp2* in the corresponding female offspring. Data are mean ± SEM; each point represents one animal (one male or female per litter; n = 12). Two-way ANOVA (factors: treatment and diet) with Tukey’s post-hoc test; *p < 0.05, **p < 0.01, ***p < 0.001; ns, not significant.

**Supplementary Figure S3. Post-diet transcriptional patterns of thermogenesis- and ETC/OXPHOS-related genes in BAT.** Bulk RNA sequencing was performed on brown adipose tissue (BAT) from F1 and F3 offspring at 18 weeks of age, after 6 weeks of high-fat diet feeding. (A–D) Volcano plots of thermogenesis-related genes in F1 (A) and F3 (C) males and electron transport chain/oxidative phosphorylation (ETC/OXPHOS)-related genes in F1 (B) and F3 (D) males, comparing TBT-line offspring with their corresponding DMSO controls. Each point represents one gene. The x-axis shows log2 fold change (TBT/DMSO), and the y-axis shows −log10(*P* value). Significantly increased and decreased genes are shown in red and blue, respectively; nonsignificant genes are shown in gray. The horizontal dashed line indicates *P* = 0.05. (E, F) Heatmaps showing post-diet TBT-to-DMSO log2 mean expression ratios for thermogenesis-related (E) and ETC/OXPHOS-related (F) gene panels in F1 and F3 male and female BAT. Blue indicates lower expression in TBT-line BAT, white indicates little or no change, and gold indicates higher expression relative to DMSO controls. After dietary challenge, expression changes in these gene panels varied by sex and generation.

**Supplementary Figure S4. Differential expression of thermogenesis- and mitochondrial respiratory-chain-related genes in pre- and post-diet female BAT.** Bulk RNA sequencing was performed on brown adipose tissue (BAT) collected from F1 and F3 female offspring at 12 weeks before high-fat diet challenge (Pre-Diet) and at 18 weeks after 6 weeks of high-fat diet feeding (Post-Diet). (A, C) Volcano plots of genes associated with the thermogenesis pathway in pre-diet F1 (A) and F3 (C) TBT-line females compared with their corresponding DMSO controls. (B, D) Corresponding volcano plots of electron transport chain and oxidative phosphorylation (ETC/OXPHOS)-related genes in pre-diet F1 (B) and F3 (D) females. (E, G) Volcano plots of thermogenesis-related genes in post-diet F1 (E) and F3 (G) females. (F, H) Corresponding volcano plots of ETC/OXPHOS-related genes in post-diet F1 (F) and F3 (H) females. Each point represents one gene. The x-axis shows the log2 fold change in TBT-line relative to DMSO control BAT, and the y-axis shows −log10(*p* value). Significantly increased and decreased genes in TBT-line BAT are shown in red and blue, respectively, while nonsignificant genes are shown in gray. Top 10 changed genes are labeled, and the horizontal dashed line indicates *p* = 0.05. Female BAT exhibited gene-, generation-, and diet-dependent changes in both thermogenesis- and ETC/OXPHOS-related transcriptional programs, without a consistent direction of regulation across groups.

**Supplementary Figure S5. Relative mitochondrial DNA content in male and female offspring and electron transport chain respiratory capacity in pre-diet female BAT.** (A–B) Relative mitochondrial 16S rRNA and ND1 copy numbers in brown adipose tissue (BAT) from F1 and F3 male offspring at 12 weeks (Pre-Diet). (C–D) Corresponding measurements in F1 and F3 female offspring. No consistent differences in relative mtDNA content were observed between DMSO- and TBT-line offspring. No significant differences were detected in males or in female ND1 copy number, although 16S copy number was increased in F1 TBT-line females. (E–F) Oxygen consumption rate (OCR) profiles obtained using the electron transport chain protocol (ETCP) in F1 and F3 female pre-diet BAT, respectively, following sequential addition of rotenone/succinate, antimycin A, ascorbate/TMPD, and sodium azide. (G–I) Quantification of complex I-, complex II-, and complex IV-associated respiratory capacity in F1 female BAT. (J–L) Corresponding quantification in F3 female BAT. No significant differences in complex I, complex II, or complex IV respiratory capacity were detected between DMSO- and TBT-line females in either generation. All values are shown as mean ± SEM. Data were analyzed by unpaired *t*-test. *p < 0.05; ns, not significant.

